## Supplementary material for "Coupling of Lipid Phase Behavior and Protein Oligomerization in a Lattice Model of Raft Membranes": SI Figure

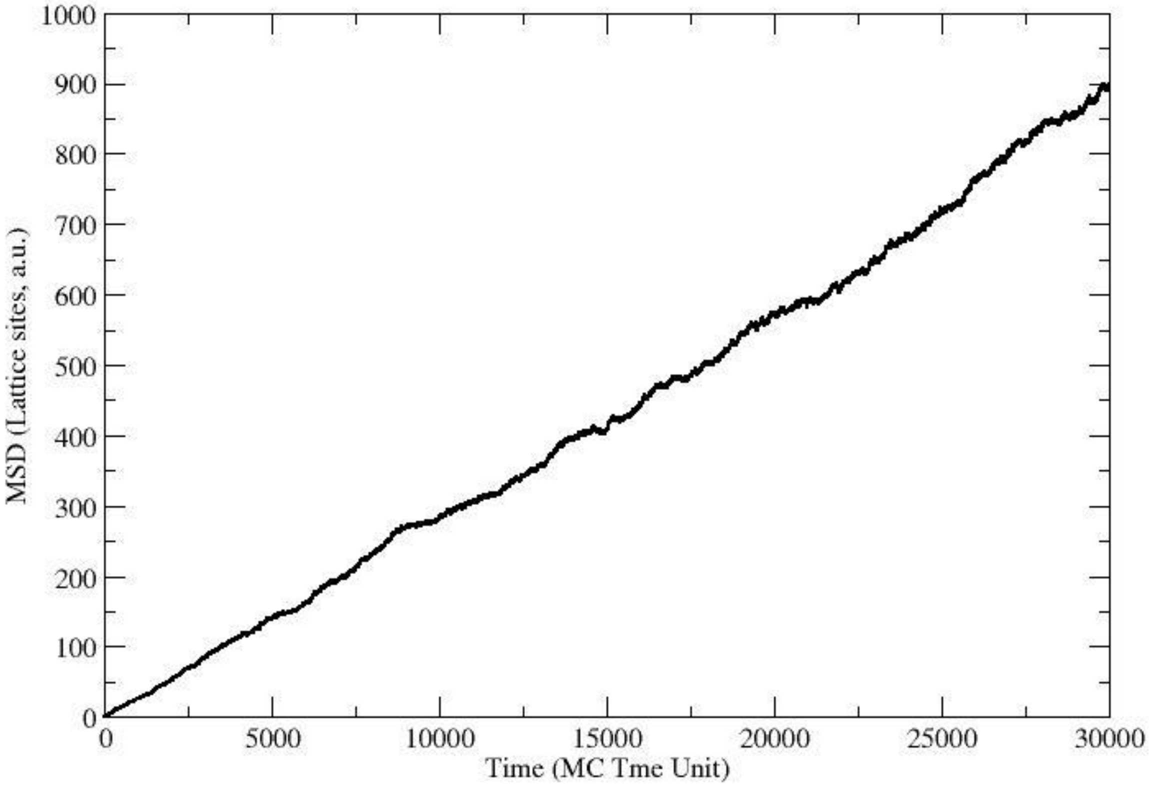


**Fig. S1**: Mean square displacement of a single protein in a 35DPPC tenrary mixture.
